# The ParA’s function is realized by two separate proteins in the partitioning system of *Myxococcus* plasmid pMF1

**DOI:** 10.1101/2020.07.24.219303

**Authors:** Duohong Sheng, Xiaojing Chen, Yajie Li, Jingjing Wang, Li Zhuo, Yuezhong Li

## Abstract

The *par* operon in the sole myxobacterial plasmid pMF1 includes a function-unknown *parC* gene in front of the classical *parA* and *parB* genes. Removal of *parC* severely reduced plasmid stability, but ex-situ compensations of *parC* did not restore the *par* system function. Individual expression of *parA* formed insoluble proteins, while co-expression of *parC* before *parA* produced a soluble ParC-ParA heterodimer. ParA alone had no ATPase activity and no polymerization, while ParC addition aided ParA to restore the activities. Fusing ParC and ParA in different ways all produced soluble proteins and some restored ATPase activity or increased plasmid stability. Protein interaction model analysis and experiments revealed that ParC structurally mimics the N-terminal of Ia-type SopA (ParA), endowing the *Myxococcus* ParA protein to play functions by shifting of ParC between two sites on ParA surface. The present results highlight that ParC functions as a part of ParA to support its soluble expression and function, and the separation of ParC and ParA into two proteins in structure enables the ParC ‘fragment’ to shift in a larger range around ParA to function during partitioning.

**Author summary:** Our work on ParC here provides a new example for the evolution of multi-domain protein. ParC and ParA are two proteins, but their expression and function act as a whole, which proposes a new regulatory model for bacterial *par* system, and also provides research ideas and materials for the study of functional coordination and evolution of ParA domains in the future.

## Introduction

Partitioning (*par*) system exists widely in bacteria and archaea, participating in the isolation and allocation of chromosome and plasmid into daughter cells during the cell division [1–3]. A *par* system typically consists of three components: an NTPase (usually named ParA), a DNA-binding protein ParB, and one or more *cis*-acting sequences *parS*. ParB binds specifically onto the *parS* sequences and assembles into a high-order partitioning complex, on which the ParA proteins are further added by binding to the ParB proteins, and act as an energy machinery by hydrolyzing NTP to move the ParB-*parS* complex apart. The two *par* genes are usually found in the same operon, with the *parS* elements located within or adjacent to this operon [4–7].

According to the ParA sequence similarity, bacterial *par* systems are divided into three main types [7,8]. Type I *par* system, which has a deviant Walker-box ATPase, is found in most bacterial and archaeal chromosomes and plasmids, and is probably the most ubiquitous type of active partitioning system in nature. In addition to ATPase activity, Type I ParA has the capability of auto-aggregating into dimers and then more complex polymers [9–11]. Type I *par* system can be further divided into two subfamilies, Ia and Ib [12–14]. Type Ia ParA, such as the ParA proteins encoded in P1 and F plasmids, contains a Walker-box region and an N-terminal winged helix–turn–helix (HTH) domain, which plays an auto-regulation role for the transcription of *parAB* operon [11–14]. Type Ib ParA is some small proteins (192–308aa), containing only the Walker-box domain, represented as the ParA proteins in pTAR, TP228, pB171 or pSM19035, and their auto-regulation role for *Par* transcription is fulfilled by the ParB proteins [7,8,15]. HTH-containing and -free ParAs both use the Walker-box domain to engage in the nucleoid to track for their partitioning functions [16].

The pMF1 plasmid was discovered in *Myxococcus fulvus* 124B02, and is the first and as yet the sole endogenous plasmid that is able to replicate autonomously in myxobacterial cells [17]. The plasmid is 18,634 bp in size, containing 23 genes (*pMF1.1-pMF1.23*). Many of the plasmid genes have their homologs in different myxobacterial genomes, while the others have not yet found their homologs in the GenBank database (**S1 Table**), which suggest that pMF1 has a long standing co-adaption within myxobacteria [18,19]. We previously determined that the plasmid replication is controlled by the *pMF1.13-pMF1.15* fragment (gene locations refer to **S1A Fig**), based on which, shuttle vectors between *Escherichia coli* and *M. xanthus* DZ1 have been successfully constructed [17,20,21]. pMF1 employed at least two strategies for its stable inheritance in *Myxococcus*; one is the partitioning system encoded by the *pMF1.21-pMF1.23* genes [22], while the second is a post-segregational killing system of nuclease toxin and immune protein encoded by the *pMF1.20* and *pMF1.19* gene pair [23]. In the partitioning system, *pMF1.22* is predicted a *parA* gene, while*pMF1.23* encodes a DNA-binding protein (*parB*), which is able to specifically bind to the *parS* iteron DNA sequences [22]. Based on the ParA protein sequence similarity and the autogenously repressing regulation of ParB on the expression of the *par* genes, the pMF1 partitioning system was suggested to belong to the Ib system [22]. Specifically, the pMF1 *par* system has a third function-unknown gene, *pMF1.21* (named *parC*), in front of *parA* and *parB*. The three genes are in the same transcriptional unit, and *parC* is able to enhance the DNA-binding activity of ParB and regulate gene expression of the *par* loci [18,22,23].

In this study, we revealed that *pMF1.21* is functionally essential for the partitioning system in *Myxococcus* cells. We determined that *parC* located in front of *parA* is critical for regulating the expression and function of ParA. There are two ParC binding sites on the surface of ParA protein, C1 and C2. Shift of ParC between the binding regions of ParA affects the expression and function of ParA. This novel molecular mechanism for plasmid partitioning not only provides a clue for explaining the polymer filament formation of the Walker-box ParA proteins, but also implies a new potential approach in protein engineering.

## Results

### 1. *parC* and its location in the *par* system are critical for plasmid partitioning

ParC is a small acidic protein with 87 amino acid residues and the *parC* gene locates in front of *parA* in the pMF1 partitioning operon. Quantitative PCR amplification indicated that, in the pMF1-harboring *M. fulvus* 124B02 strain, as well as the *par* system-containing shuttle plasmid pZJY4111, *parC* transcribed slightly higher than *parA*, but lower than *parB*, which were comparatively at similar transcriptional levels as the replication genes (**S1B Fig**). To determine whether *parC* is required for the partitioning function, we made an in-frame deletion of the *parC* gene in pZJY4111 (**Fig 1A**), which contains the *ori* region and *par* system of pMF1 [22]. Compared to approximately 60% retention of pZJY4111 in DZ1 strain after 144 h of incubation in the absence of antibiotic selection, the retention of pZJY4111Δ*parC* was dramatically declined to 20% after 48 h, and to almost zero after 96 h of incubation (**Fig 1B**). The above results indicated that the *parC* gene plays a crucial role in the partitioning system for the plasmid maintenance in *Myxococcus* cells.

**Figure 1.**
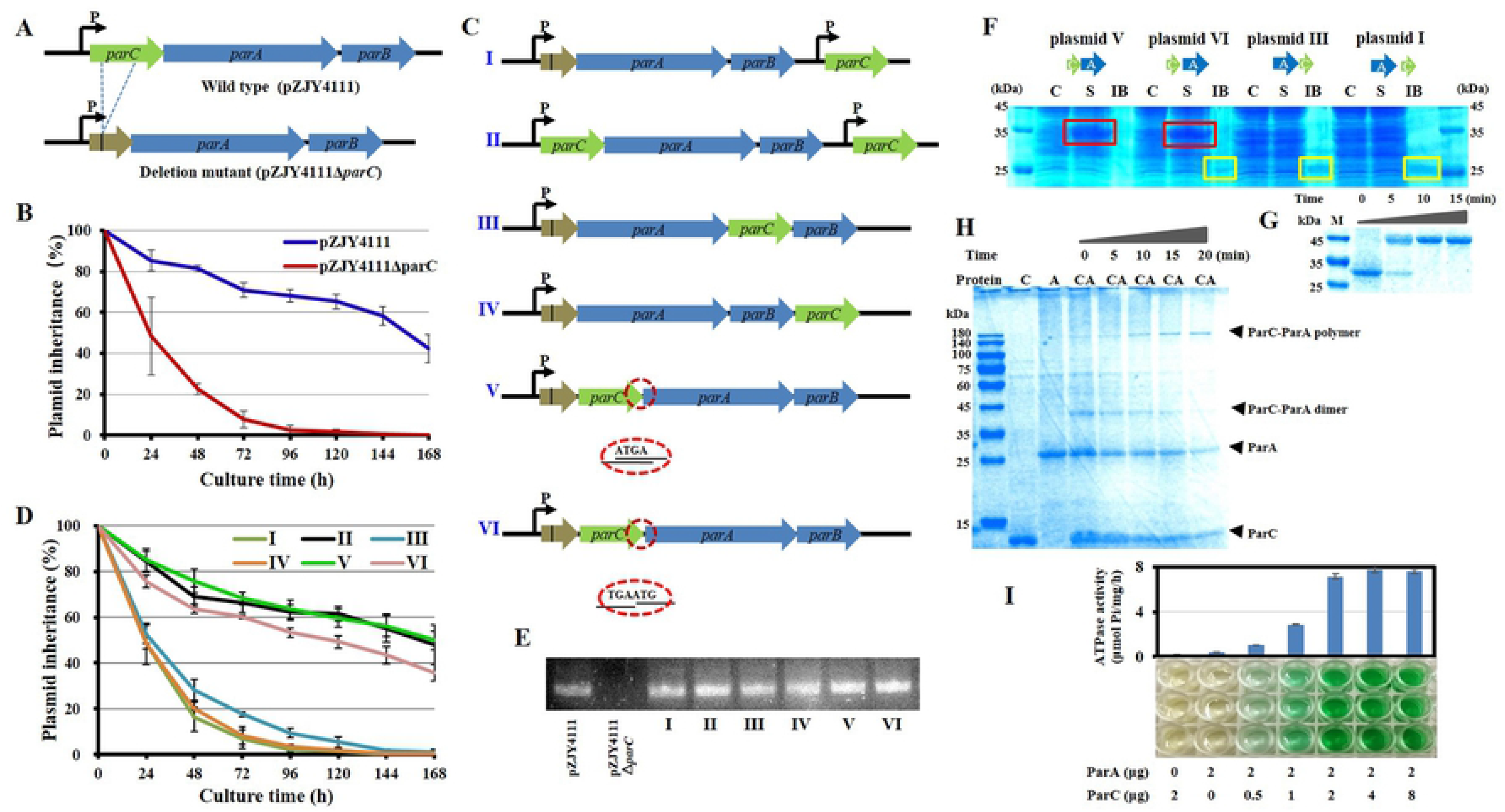
Mutation and organization of the *parC* gene in the shuttle plasmid pZJY4111 and its effects on the inheritance stability of plasmid. (**A**) In-frame deletion of *parC* of the *parCAB* operon in pZJY4111. The deletion removed 228 bases and retaining 11 N-terminal bases and 25 C-terminal bases of *parC* (**S8A Fig** in **Supplementary materials**), producing plasmid pZJY4111Δ*parC*, which was further electro-transformed into *M. xanthus* DZ1 to assay the plasmid retention ability. Arrows indicate the direction of transcription. (**B**) Inheritance stability of pZJY4111 and pZJY4111Δ*parC* in *M. xanthus* DZ1. (**C**) Schematic representation for different locations of the compensatory *parC* gene in pZJY4111Δ*parC* or pZJY4111. These plasmids were separately transformed into *M. xanthus* DZ1 to assay their inheritance stabilities in *M. xanthus* DZ1 (**D**). Error bars in **B** and **D** indicate the standard deviation (SD) of at least five independent experiments. The DZ1 strains with different plasmids were cultivated in CTT medium without antibiotics for 168 h, and the plasmid maintenance were examined every 24 h. (**E**) Transcriptional confirmation of the expression of *parC* in *M. xanthus* DZ1 containing different plasmids using quantitative RT-PCR. (**F**) Soluble expression analysis of ParC and ParA proteins in *E. coli* in different arrangements of *parC* and *parA* (the original gel picture is provided as **S8B Fig** in **Supplementary materials**). The recombination plasmids were transformed into *E. coli* BL21(DE3) and expressed by IPTG induction. After harvested, cells were washed, broken and centrifuged. The supernatant (S) and inclusion body (IB) were analyzed by SDS-PAGE. To aid proteins dissolve for electrophoresis, IB was homogenized using tissue grinder and treated with 6M guanidine hydrochloride (GuCl) for protein dissolution. Non-induced cultures were used as control (C). Red box frames the induced soluble proteins (35kD), while yellow box frames the insoluble proteins with the ParA size. (**G**) ParA monomer was incubated at 32 °C and sampled at intervals (0, 5, 10 and 15 min), and the aggregation was mixed with 0.1% SDS and detected by native-page electrophoresis (the original gel picture is provided as **S8C Fig** in **Supplementary materials**). (**H**) Interactions between ParA and ParC in binding buffer containing 3mM ATP, incubated at 32°C. The mixture was sampled at intervals of 0, 5, 10, 15 and 20 min, and detected by electrophoresis. (**I**) ATPase activity assay with ParC and ParA proteins. ATPase activity was measured by detecting the release of Pi at 32 °C. Data shown are averages of three repeats. Error bars refer to the SD.

We experimentally compensated the deletion by ex-situ inserting the *parC* gene with its own promoter in pZJY4111Δ*parC*, producing pZJY4111Δ*parC*::*parC* (plasmid I in **Fig 1C**). We performed the same insertion of the *parC* gene with its own promoter in the pZJY4111 plasmid, producing pZJY4111::*parC* (plasmid II). There was a 203-bp interval space sequence between the deficient *par* operon and the *parC* compensation gene. However, the compensation did not restore the plasmid stability phenotype, and plasmid II exhibited similar inheritance ability as pZJY4111 in *M. xanthus* DZ1 (**Fig 1D**). To verify the location effects of *parC* on plasmid stability, we placed *parC* at different places in the deficient *par* operon of pZJY4111 Δ*parC*, either after *parA*, forming the *parA-parC-parB* cascade (plasmid III); after *parB*, forming the *parA-parB-parC* cascade (plasmid IV); or in front of *parA* and behind the *parC* residuals (plasmid V) (**Fig 1C**). The plasmid stability assays indicated that plasmid III or IV did not restore the plasmid maintenance; whereas plasmid V completely restored the plasmid stability with almost the same curve as the wild type plasmid pZJY4111 (**Fig 1D**).

Notably, in the original *parCAB* operon, the coding sequences of *parC* and *parA* overlaps four bases (ATGA), and the inserted *parC* gene in plasmid V also overlapped the four bases with the *parA* gene. To investigate whether the overlap had an effect on partitioning, we added the *parC* gene in front of *parA* by replacing ATGA with TGAATG (plasmid VI; **Fig 1C**). Stability of plasmid VI reduced to some extent as compared with pZJY4111or in-situ compensating plasmid V, but significantly higher than those plasmids with ectopically inserted *parC* (**Fig 1D**). The transcriptions of *parC* in *M. xanthus* DZ1 were confirmed by quantitative PCR amplification (**Fig 1E**). The above results indicated that ectopically expressed ParC or excessive *parC* gene has no effect on the plasmid stability. The function of ParC for the plasmid partitioning strictly depends upon its adjacent location to the *parA* gene. The co-expression schedule of *parC* and *parA* was probably for interaction or functional collaboration of two proteins.

### 2. *parC* in front of *parA* promotes soluble expression of ParA by forming ParC-ParA heterodimer to play functions

To investigate the interactions between *parA* and *parC* in expression, we constructed the two genes into the pET15b expression vector in the same arrangement as they were in the compensation plasmids, and expressed them in *E. coli*, respectively. When *parC* and *parA* overlapped four bases (as in plasmid V), an obvious soluble band with approximately 35 kDa in size was induced (**Fig 1F**). For the adjacent*parC* and *parA* genes without overlapped bases (as in plasmid VI), the same soluble protein band was observed, but with a small amount of insoluble 24 kDa proteins (in ParA size). However, when *parC* was placed behind *parA* (as in plasmid III) or the two genes were transcribed separately (as in plasmid I), no obviously soluble desired band appeared. The results indicated that when *parC* was in front of *parA*, the expressed ParC and ParA proteins appeared to combine immediately into soluble ParC-ParA heterodimer; otherwise, they were in insoluble forms, probably as monomers or oligomers.

The 35 kDa band in Figure 1F was cut, digested with trypsin and identified by mass spectrometry (**S2 Fig**). The mass-charge ratio (m/z) data of peptide segments were retrieved against *M. fulvus* 124b02 database and they were identified as two proteins, ParA (pMF1.22) and ParC (pMF1.21). The relative abundance of the peak area of peptide segments were quantified and the sum of all the peak area of ParA or ParC calculated ((**S2B Fig**). The coverage ratios of the identified peptide segments were 63% and 63.22% for ParA and ParC. The relative abundance of total peak area of ParA was 2.686 times that of ParC, approximately equaling to the amino-acid ratio of the two proteins (2.609). Thus, the molar ratio of ParC (8.9 kDa) and ParA (24.07 kDa) in the 35 kDa band is one to one and the exact molecular weight of this band should be 32.97 kDa.

To assay the binding power between ParC and ParA, we fused a His-tag at the N-terminus of ParC protein for purification with Ni-NTA agarose, and co-expressed *parA* and the *parC* gene in *E. coli* cells. The induced ParC and ParA proteins appeared in the supernatant fraction as a soluble heterodimer complex under the 1% SDS condition. With the concentration increase of SDS, the ParC-ParA heterodimer complex disaggregated gradually, and mostly disappeared with the treatment of 6% SDS, accompanying with the presence of ParC and ParA monomers (**S3E Fig**).

ParC was solubly expressed easily, while ParA alone was hard to obtain soluble form. After many attempts, soluble expression of ParA in *E. coli* was obtained with the MBP fusion expression system, in which a maltose-binding protein was added at the N-terminal of ParA (**S3A-C Figs**). We removed the MBP tag from the MBP-ParA fusion protein by the proteinase Factor Xa digestion (**S6 Fig**). The prepared ParA proteins were mostly in monomer form, and turned into dimer form with almost no monomer within 10 minutes at 32°C, (**Fig 1G**). SDS treatment did not dissolve the dimer, even with 6% SDS and the addition of DTT, or 6M guanidine hydrochloride (**S3D Fig**). Accordingly, co-expressed ParC and ParA proteins in proper organization were able to combine immediately to form soluble and tightly-bound ParC-ParA heterodimer complex, preventing the formation of indissoluble ParA homodimer. Because monomeric ParA proteins easily aggregated into dimers, in the following *in vitro* experiments, ParA monomer from MBP-ParA fusion protein was used immediately after digestion, separation and purification.

ParC and ParA monomers were mixed and incubated at 32°C for the aggregation assay. With the increase of incubation time at 32°C, ParC and ParA monomers gradually decreased, ParC-ParA dimer appeared first and then gradually decreased, and a high molecular weight band (greater than 180 kDa), which was identified by mass spectrometry as the polymer of ParC and ParA and their ratio is 1:1, appeared and increased (**Fig 1F**). Thus, the presence of ParC did promote the aggregation of ParA protein into ParC-ParA polymers. Further ATPase experiments found that ParA proteins alone had almost no ATPase activity, but retrieved ATPase activity when mixed with ParC (**Fig 1I**). The ATPase activity reached the highest at the ParC-ParA ratio of 1:1, and more ParC had no increase to the activity.

### 3. Structural interactions between ParC and ParA proteins

To investigate the potential interaction patterns between ParC and ParA, we constructed their structure models using the i-tasser (Iterative thread assembly optimization) method [27]. The *Myxococcus* ParA protein was structurally composed of five beta folds in center and outside around with eight alpha helixes (**Fig 2A**). Compared with the typical ParA proteins, the *Myxococcus* ParA protein is incomplete in the N-terminal region, for example, lacking the H3-β3-β4-H4 structure when compared with Soj [25], or lacking H3 when compared with ParF [26] (**S4 Fig**). ParC, which has no significant homology with any known proteins [22], consists of three helixes in its secondary structure, and approximately 80% amino acids in the protein participate in the formation of the alpha-helix. Interestingly, the ParC secondary structure is highly similar to that of N-terminal of Ia-type ParA (SopA from plasmid F) or C-terminal of ParB (**S5 Fig**), both of which were reported to be related to the regulation of *par* operon [7,8]. ParC was also reported to involve in *par* regulation [22].

**Figure 2.**
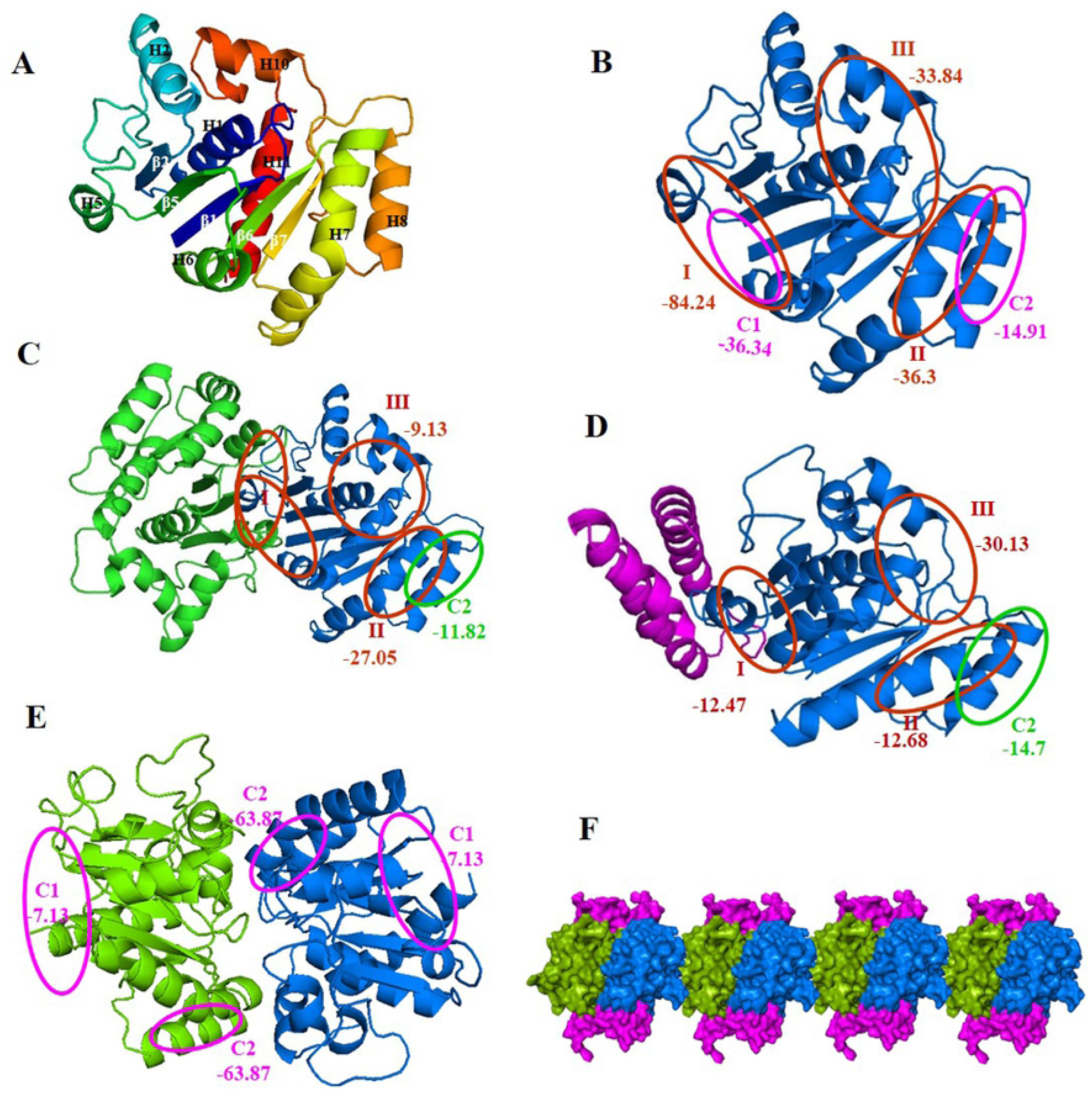
Structural basis for the protein interaction of ParA and ParC, and effects of the binding of ParC on the ParA dimerization. (**A**) Structure of ParA, predicted with I-TASSER (https://zhanglab.ccmb.med.umich.edu/I-TASSER) [27]. Myxococcal ParA structure contains β1(residues 1–6) - H1(residues14-27) - β2(residues31-37) - H2(residues42-46) – H5(residues70-71,75-77) - β5(residues82-87)-H6(residues93-101) - β6(residues103-109) – H7(residues114-115,119-128) - β7(residues137-143) – H8(residues151-161) – H10(residues176-183,187-190) – H11(residues195-213). The arrangement order of sheets and helices follows Soj. (**B**) Predicted protein-protein interaction sites on the surface of ParA. Three ParA self-dimerization regions (I, II, II) and two ParC-binding regions (C1 and C2) are marked with red and green rings, respectively. The binding energy was calculated by PRISM webserver. (**C**) Predicted protein-protein interaction sites of ParA dimer in I region (ParA-(I)-ParA). (**D**) Predicted protein-protein interaction sites on the surface of dimer of two ParC-(C1)-ParA heterodimer. (**E**) Predicted protein-protein interaction sites on the surface of dimer of two ParC-(C2)-ParA heterodimer. (**F**) Construction of ParC-(C2)-ParA polymer. ParC colored with purple red. Two paired ParAs are colored rainbow in the cartoon model and blue and green in the schematic diagram respectively.

Predicted with the PRISM web server [28], there are three potential surface regions for the formation of ParA homodimer with the binding energies of −84.24, −36.3 and − 33.84 kJ/mol, respectively (labeled as I, II and III in **Fig 2B**). Region III is similar to the dimer binding interface of ParF or Soj, which involves in the formation of P-Loop, forming an active nucleotide “sandwich” dimer with ATP in the middle [24–26]. Region II is on the outside of H7/8, partly overlapped with the binding interface 2 of ParF. Region I locates in the H5-β5-H6 region, covering an irregular sequence in front of H5. Region I has the highest binding energy, far exceeding the other two sites, leading to the preference of binding upon region I to form ParA homodimer (ParA-(I)-ParA). However, homo-dimerization on region I reduces the binding ability of the other two aggregation sites, especially in region III, the binding energy of which turns to −9.13 kJ/mol in the homo-dimer (**Fig 2C**), hindering further active dimerization at region III, which might explain why ParA alone had no ATPase activity and was not further polymerized.

Besides the three homo-dimerization regions, two regions on the surface of ParA were predicted for ParC binding, with −36.34 kJ/mol and −14.91 kJ/mol binding energy, respectively (labeled C1 and C2 in **Fig 2B**). C1 locates in the N-terminal region, and covers the region of H5, H5-β5 loop, H6-β6 loop and C-terminus of H11, which partially overlaps with region I. C2 and the self-dimerization region II are both located on H7/8 with some overlaps. C1 is the main binding site of monomers ParA and ParC. Binding of ParA with ParC significantly decreases the self-dimerization binding energy of region I and II, from −84.24 and −36.3 kJ/mol to −12.47 and −12.68 kJ/mol kJ/Mol, respectively (**Fig 2D**). Thus, binding with ParC results in region III becoming the main ParA self-dimerization site in the ParC-ParA heterodimer complex, which might be the reason why ParC-ParA has ATPase activity.

The ParC-A-A-C tetramers formed on region III have the ability to further aggregate on region II in theory. However, dimerization of ParA at site III affects ParA’s 3D structure and the binding abilities between ParA and ParC decreases from − 36.34 to −7.13 at C1 site, and increases significantly from −14.91 to −63.87 at C2 site, respectively (**Fig 2E**). As a result, ParC on ParA dimer in the ParC-A-A-C tetramer moves from C1 to C2 site, and the site of the tetramer for further aggregation is changed from region II to region I (**Fig 2F**).

### 4. ParC locating shift on C1 and C2 sites of ParA for soluble expression and plasmid stability

In the structural model of ParCA, the ParC fragment was near the C1 binding site (**S5C Fig**). Region I is affected, forming a new N-terminal region for self-dimerization binding (−12.47 kJ/Mol). Similar to that in ParA, the ParCA fusion protein retains the II and III self-aggregation regions, but with significantly decreased binding energy of region II (−12.68 kJ/mol). It was speculated that the ParCA protein preferentially forms homodimer at site III and thus has ATPase activity. To test the hypothesis, we fused the ParC and ParA proteins by adding an A base before the overlapping area (ATGA) of *parC* and *parA* to destroy the stop codon of *parC* (TGA) (plasmid VII). Compared to the wild-type pZJY4111 plasmid, plasmid VII had a much low retention capacity in *M. xanthus* DZ1, which was at the similar level as the *parC* deletion mutant. The ParCA fusion protein was solubly expressed in *E. coli*, together with some inclusion bodies. ParCA exhibited ATPase activity, which, however, was much lower than that of ParC-ParA heterodimer (**Fig 3A**). Because the binding energies of the N-terminal binding region and region II in the ParCA-ParCA dimer (binding at region III) were similarly low, there were two polymerization patterns via the N-terminal or II site (**S5D Fig**). Similar and low binding abilities resulted in low efficient binding competition, which might be the reason for the low activity of the ParCA fusion protein.

**Figure 3.**
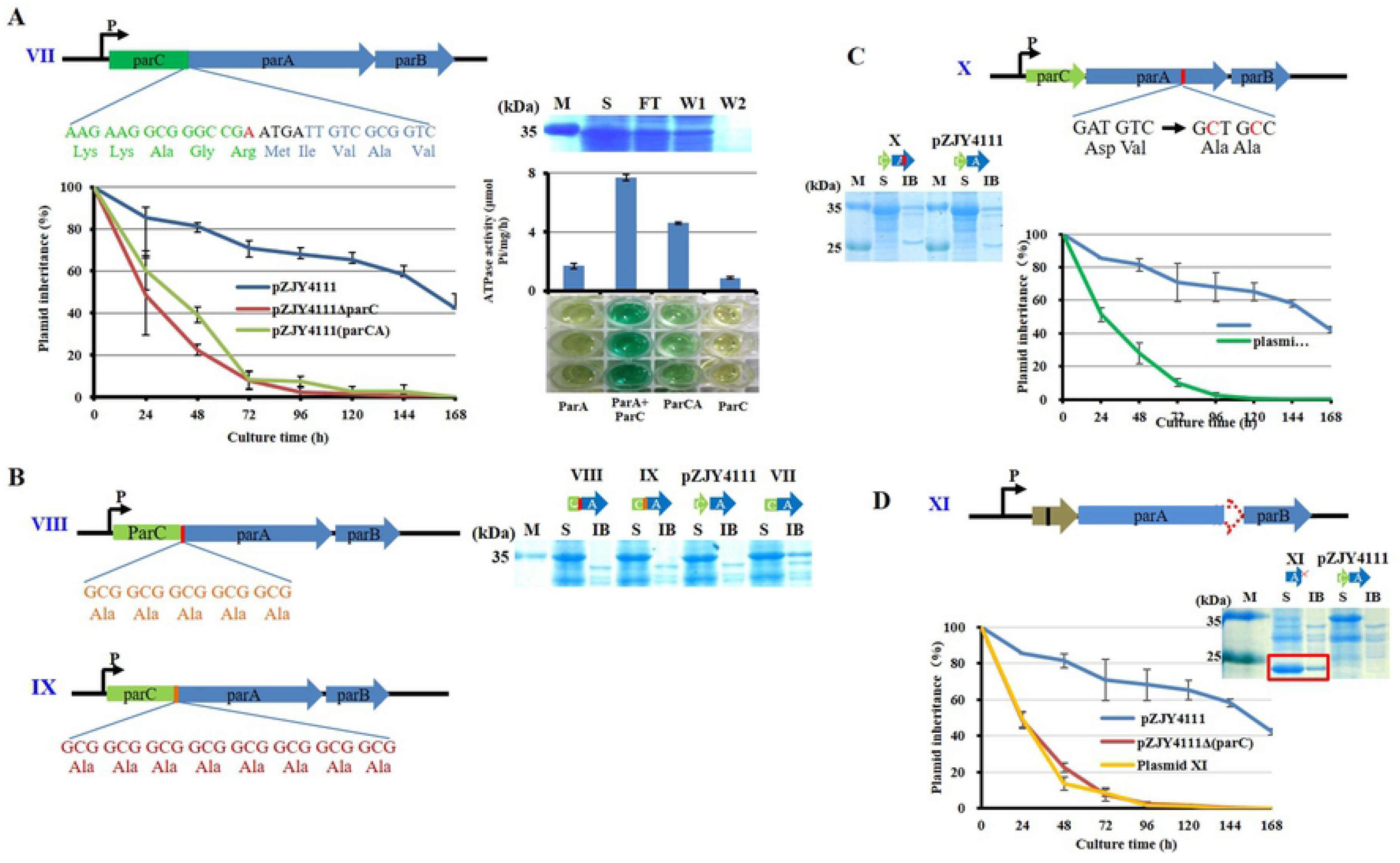
Properties of ParCA fusion proteins. (**A**) An adenylate was inserted before the overlapping region (ATGA) to destruct the stop codon (TGA) of *parC*, forming a fused ParCA (plasmid VII). (**B**) Adding a 5×Ala linker (VIII) or an 8×Ala linker (IX) between domains C and A in ParCA fusion protein. (**C**) Amino acid mutation at C2 site in ParA proteins (plasmid X). (**D**) Deletion of 42 C-terminal bases (14 amino acids) of *parA* of the parCAB operon in pZJY4111Δ*parC* (plasmid XI). Inheritance stabilities of different plasmids was assayed in *M. xanthus* DZ1. Error bars indicate the SD of four or six independent experiments. Expression of the ParCA fusion proteins was performed in *E. coli* BL21(DE3) cells. ATPase activity was assayed of the ParCA fusion protein. ParA, ParC and ParA+ParC proteins as control. M, protein standard; S, supernatant; FT, flowthrough Ni-NTA column; W, washing with 20 mM imidazole. Red box in (**D**) frames the target protein band in the supernatant (S) and inclusion body (IB). The original gel pictures are provided as **Figures S9** in **Supplementary materials**.

Fused ParCA protein improved soluble expression and ATPase, but did not promote plasmid stability, which is most likely due to two possible reasons: the fusion affected the flexibility of further binding activity, and/or ParA functioning in plasmid partitioning required dissociation or movement of ParC fragment from C1 site. To improve flexibility between the ParC and ParA fragments in the fusion protein, we added a five-alanine or eight-alanine hinge between them (VIII and IX in **Fig 3B**). The two fusion proteins were almost completely soluble in *E. coli*, which suggested that ParC binding to C1 region promoted the protein solubility. Interestingly, while plasmid VIII had no significant effect on plasmid stability, the plasmid IX with a longer amino acid linker improved the plasmid stability to a similar level as pZJY4111. The results suggested that fixation of ParC on the C1 region is not helpful for plasmid partitioning, which requires a larger moving range of ParC; this supports the above dynamitic structural modelling of movement of ParC on ParA from C1 to C2.

As PRISM predicted, the C2 region mainly distributed in H8, of which the key amino acids D157 and V158 were not overlapped in the ParA self-aggregation region II (**S6 Fig**). We mutated the C2 region by substituting D157V158 to A157A158. As predicted, the binding ability of the ParA mutant to ParC at C2 will decrease, and a third ParC binding site will appear at the H11 region. With the dimerization of ParA with C2 site mutation, ParC did not bind to C2 site, but to H11 helix (**S6 Fig**). Experimental assays showed that the point mutations in C2 region did not affect the soluble co-expression of ParC / ParA protein, but significantly reduced the retention of plasmid X (**Fig 3C**), which further confirmed that the binding of ParC and ParA at C2 region is important for the plasmid partitioning.

In typical *par* systems, ParB is usually an alkaline protein, while ParA is an acidic protein. However, in our pMF1 *par* system, ParA is also an alkaline protein (the theoretical pI is 8.90), which is mainly attributed by the rich basic amino acids in the C-terminal (**S4 Fig**). Interestingly, ParC is an acidic protein (pI is 4.57), and the theoretical pI of the ParC-ParA heterodimer is 5.68. The C-terminal and the N-terminal are close in structure of the known Ib-type ParA. ParC is predicted to locate at N-terminal, which probably favors for charge neutralization. We prepared the C-terminal deletion mutant of ParA by removing 14 C-terminal amino acids. Without the basic tail, the ParA mutant proteins were solubly expressed in *E. coli*, but had no distribution for the plasmid stability in *Myxococcus* cells (**3D Fig**). The C-terminal alkaline tail of ParA may be related to its improper folding or insoluble expression.

## Discussion

pMF1 ParA has the typical structure of Walker ATPase protein without the N-terminal HTH domain, and belongs to the ParA Ib family, similar to the ParF of *Salmonella newport* TP228 plasmid and Soj of *Thermus thermophilus* [24–26]. However, compared with typical Ib ParA proteins [25,26], pMF1 ParA is incomplete. ParA alone cannot fold correctly and exhibits no ATP activity. The protein binds tightly to form homodimer, preventing further polymerization. Here, fore-expressed ParC interacts immediately with new-born ParA at a ratio of 1:1, forming an active heterodimer to prevent the ParA homo-dimerization, and thus participates in the process of plasmid partitioning. ParC unlocks activities that ParA deserves, and seems to be a subunit of a complete ParA protein for function. ParC has no homologous protein in GenBank, but is structurally similar to the N-terminal fragment of type Ia ParA of plasmid P1 [12,29] or the C-terminal fragment of *Sulfolobus* ParB [30], both of which involve in the regulation of *par* operon [22,29–31]. ParC binds to ParA on the C1 binding site near the N-terminus (**Fig 4B**), mimicking the N-terminal fragment of the Ia ParA protein.

**Figure 4.**
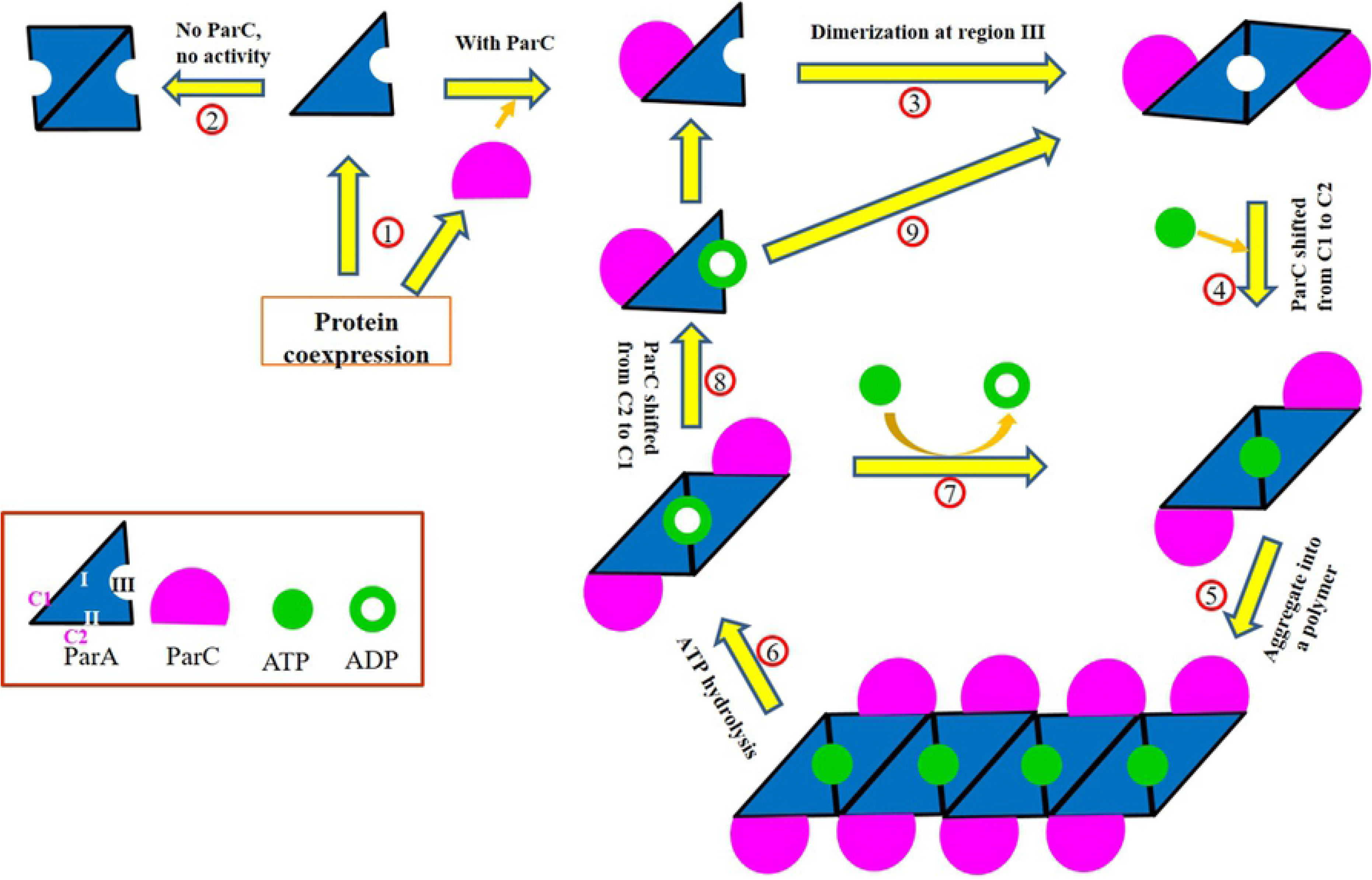
A possible model of ParC regulating the expression and function of ParA. Coexpression of ParC and ParA to form soluble binding complex 1. ParA alone tends to form inactive dimer at I region2. The dimer formed by ParC and ParA at C1 site is further homodimerized in region III to form ATP loop3. Then, ParC shifted from C1 to C24 and ATP combining to ParA promotes further aggregation of CAAC tetramer5. With ParA playing its role in plasmid partition, 6ATP is hydrolyzed to ADP, and the polymer is depolymerized. The depolymerized CAAC tetramer with ADP, which can be replaced by ATP to re-enter the polymer cycle7, or further depolymerized into CA dimer8. In the CA dimer, ParC moves back to the C1 site. Then CA further dimerization 9, once again into the PA aggregation cycle.

Polymerization or depolymerization of ParA are related to the ATP binding, hydrolysis and ADP replacement [12–15]. Calculation of the binding abilities (**S7 Fig**) shows that when ParA was dimerized, whether ParA bound ATP or ADP, ParC is only able to bind to C2 site. When ParA was in the monomer form, ParA-ADP binds to ParC at C1 site, but ParA-ATP binds to ParC at both C1 or C2 sites and preferentially at C2 site. ParA-ATP binding ParC at C1 site forms an active dimer in III region, but forms an inactive dimer when binding ParC at C2 site. This means that monomeric ParA-ATP binding ParC at C2 site has no contribution to activity or dimerization of ParA; ParA is functional to form dimers only when binding ParC at C1. Thus, ParC needs to move between C1 and C2 sites in the ParA functioning process. Firstly, ParC binds ParA monomer at C1 to stabilize protein structure. At the same time, ATP is added to promote the further aggregation of the heterodimers at the N-terminal (I). Then, ATP is hydrolyzed into ADP, ParA polymer is depolymerized to dimer or monomer in which ParC moves back to C1 position (**Fig 4**).

Gene fusion/fission is a major contributor to evolution of multi-domain proteins [42]. Here, ParC and ParA function as a whole, but separate into two proteins, which provides more interaction plasticity and thus is more efficient than one fusion protein in plasmid partitioning. Their interfaces can be changed in a larger range, which contributes to better protein–protein interactions and increases the protein activity and precision adjustability. This is a novel regulation method of bacterial *par* system, different from any known Ia or Ib ParA proteins and will be a good model for studying the function of type I ParA [32].

## Materials and methods

### Strains, plasmids, and culture conditions

The strains and plasmids used in this study are listed in **S2 Table**. The *M. fulvus* 124B02 and its derived strains were cultivated routinely in CTT medium at 30 °C [35]. The *Escherichia coli* strains were cultivated in LB medium at 37°C. When required, 40 μg/ml of kanamycin and 100 μg/ml ampicillin were added into the medium.

### Plasmid curing

The principle that the plasmids with the same *ori* region are incompatible was employed to cure pMF1 [36,37]. The plasmids of pZJY4111 [22] and pMF1 shared the same *ori* and *par* loci. The pMF1 curing in *M. fulvus* 124B02 was conducted using the protocol as previously described with some modifications [38]. The shuttling plasmid pZJY4111 was introduced into *M. fulvus124B02* with pMF1 by electroporation (400 Ω, 25 μF, 1.25 kV). Then, the cells were grown at 30°C for 4 h in CTT broth without antibiotics on a shaker (200 rpm), followed by spreading onto kanamycin-containing (40 μg/ml) CTT agar plates. After 5 to 6-dayincubation, resistant clones were retransferred onto kanamycin-containing agar plates. Then three clones were selected, scattered by magnetic beads, and incubated in 50 mL kanamycin-containing CTT broth (30°C, 200 rpm) for 24 h. After seven times of transfer, aliquots of each culture were diluted and spread onto kanamycin-containing agar plates. To determine the elimination of pMF1 and presence of pZJY4111, the total genomic DNA of some clones was extracted and analyzed by PCR amplification of different regions. The primers used in this study are listed in **S3 Table**. Positive clones were selected and designated as *M. fulvus* 124B02/pZJY4111.

Next, the 124B02/pZJY4111 was cultured in 50 ml CTT broth without antibiotics, transferred as above for seven times, and spread on plates without antibiotics. After 6day incubation, single clones were transferred onto plates with and without kanamycin to screen plasmid-free candidate.

### Construction of *parC* knockout, compensation and overexpression plasmids

Based on the pZJY4111 plasmid, the deletion, compensation and overexpression plasmids of *parC* gene were constructed by PCR and DNA ligation. The used primers were listed in **S3 Table**. These plasmids were introduced into *M. xanthus* DZ1 by electroporation for further plasmid stability assay.

### Plasmid stability assay

To test the stability, *M. xanthus* DZ1 strains harboring the plasmids were grown to the late exponential phase in liquid CTT medium supplemented with 40 μg/ml kanamycin. Then we diluted the cultures by 1:25 in fresh CTT liquid medium with no antibiotics and grown at 30 °C and 200 rpm. After 24 h of incubation, the cultures were serially diluted and plated on CTT agar without antibiotics. The dilutions and plating were routinely repeated every 24 h till 168 h of incubation. In each round, 100 single colonies were patched onto CTT agar with and without kanamycin, and the plasmid stability was measured as the percentage of antibiotic-resistant clones [22].

### Protein expression and purification

The proteins were expressed in *E. coli* BL21 (DE3), induced by the addition of 0.1 mM of IPTG when the OD_600_ value of the culture reached 1.0. The BL21 cells were grown at 37 °C in LB broth with antibiotics. After the addition of IPTG, the cultures were grown at 16 °C for 20 h. The cells were then collected and resuspended in lysis buffer (25 mM Tris-HCl, pH 8.0, 200 mM NaCl and 5% glycerol, pH 8.0) and lysed via ultra-sonication. The mixtures were centrifuged at 4 °C for 30 min, 12000 rpm. The soluble proteins were mixed with amylose resin (New England Biolabs) according to the manufacturer’s protocols.

### Structural modeling

I-TASSER was used to model the structure of ParA and ParC based on a threading approach [27]. I-TASSER is a hierarchical method for protein structure prediction. Structural templates were first identified from the PDB by the multiple-threading program LOMETS; then, full-length models were constructed by iterative template fragment assembly simulations. All the structural models were refined in the atomic-level by the fragment-guided molecular dynamics (FG-MD) simulations [39]. The modeled structure was displayed by PyMol.

The protein structure of ParA-binding ATP/ADP was based on its homologous protein Soj with ATP/ADP (6iub/6iud), and constructed by homologous modeling method (https://swissmodel.expasy.org/).

### Protein–protein docking analysis

We further predicted the interaction pattern between the ParA and ParC proteins. The modeled structures were submitted as targets to PRISM protein–protein docking server [28]. PRISM predicts possible interactions, and how the interaction partners connect structurally, based on geometrical comparisons of the template structures and the target structures.

### ATPase Assay

The ATPase activity measurements were performed spectrophotometrically using the ultra-micro total ATPase detection kit (Jiancheng Bio. Nanjing), as previously shown [40].

### Polymerization assay

ParA polymerization assay was performed as previously described [25] with some modifications. ParA protein (6 μg) was incubated in 20 buffer (20 mM Tris·HCl pH 7.0, 100 mM KCl, 2 mM DTT, 10% glycerol, 2 mM Oligodeoxynucleotides) for 15 min at 30 °C. In ParC Co-aggregation assays, Gradient ParC was added to the reacting solution at the concentrations indicated. After the reaction completed, the samples were added 0.1% SDS, analyzed by native–PAGE and coomassie blue staining.

## Acknowledgements

This work was financially supported by the National Natural Science Foundation of China (NSFC) (Nos. 31670076 and 31471183), and the Key Program of Shandong Natural Science Foundation (No. ZR2016QZ002) to YZL.

## Supporting information

**S1 Fig**. (**A**) Organization of the genes in plasmid pMF1 and their homologous relationships with genes in myxobacterial genomes. 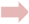, Seven hypothetical genes, each having a single homolog in *M. stipitatus* DSM 14675. 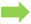, Three genes, each having many homologs in different genomes only of myxobacteria. 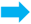, Three genes, each having a single homolog in many different genomes, but the genes with the highest similarity come from myxobacteria. 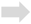, Ten genes, having no homology in the survey. Details are referred to **S1 Table**. The genes of *repA, repB, repC* (*pMF1.13-pMF1.15*), *parC, parA, parB* (*pMF1.21-pMF1.23*), *toxP* and *immP* (*pMF1.20* and *pMF1.19*) have been determined to involve in the plasmid replication, the plasmid partition, and a post-segregational killing system for plasmid stable inheritance [17, 22, 23]. (**B**) Quantitative analysis of *par* genes transcription using quantitative RT-PCR. The replication genes were inspected as a control and primers used for intergenic region are listed in **S3 Table**.

**S2 Fig**. Identification of the induced 35 kDa protein by mass spectrometry. The induced protein bands in the red box (**Figure 1F**) were cut, digested by trypsin and identified by mass spectrometry. Peptide peaks and their mass-charge spectra are shown in the above figure, and the identified proteins are listed in the following table.

**S3 Fig.** (**A**) Electrophoresis assay of the successfully soluble expressions of ParC and ParA proteins in *E. coli*. These vectors include pET15b, pET30a, pGEX-6p-1, pGro7, pKJE7, pG-Tf2, pG-KJE8, pTf16. (**B** and **C**) The induction expression of *parA* in the pMAL-c2x expression system. (**D**) Dissociation analysis of ParA homodimer with gradient SDS concentrations, combined with 10 mM DTT or 6M guanidine hydrochloride (GuCl), which was detected by electrophoresis. (**E**) Co-expression of the *parC* (fused with a His-tag fragment) and *parA* genes in *E. coli* and dissolution treatment of the purified ParC-ParA complex. The *parC* and *parA* genes were constructed in pET15b, which was electrotransformed into *E. coli* cells for expression. *The* expressed His-tagged ParC proteins was purified using Ni-NTA agarose according to the standard protocol (Qiagen, Valencia, CA). The purified proteins were treated with gradient SDS concentrations and analyzed by polyacrylamide electrophoresis. M, marker; C, control; WCP, whole cell protein; S, supernatant; IB, inclusion body; FT, flow though; W, washing; E, elution.

**S4 Fig**. Structural and sequence comparison of ParA with Soj of *T. thermophilus* [5] (**A**) or ParF of *S. newport* TP228 plasmid [26] (**B**) Protein structure was predicted with I-TASSER (https://zhanglab.ccmb.med.umich.edu/I-TASSER) [27]. In the structure comparison, ParA is shown in red and Soj or ParF is shown in blue. The differences in structure and sequence amino acids were indicated with arrows in the structure comparison.

**S5 Fig**. (**A**) Structure comparison of ParC, N-terminal of SopA, and C-terminal of ParB. N-terminal sequence of type Ia SopA is from plasmid F (PDB ID: 3EZ7.1), and C-terminal sequence of ParB is from *Sulfolobus* pNOB8 (PDB ID: 4RS7). (**B**) Multiple sequence alignment of ParF, ParA, ParCA and SopA. (**C**) Three surface self-dimerization sites are marked with red circle and their binding energy were calculated by PRISM protein-protein docking server [28]. (**D**) ParCA docking on the self-dimerization site III. Paired two ParCAs are colored rainbow in the cartoon model. Further aggregation sites of ParCA dimer have been predicted and labeled in the model.

**S6 Fig**. Mutation of the key amino acid in C2 site to destroy the binding of ParC at C2 site. (**A**) Sketch for amino-acid mutation in C2 in ParA fusion. (**B**) C2 location and amino acids involved in ParC binding. The amino acids circled by red circles are candidate amino acids for mutation. (**C**) Prediction of the interaction of ParC and ParA with amino-acid mutation in C2 site. Circled in green dotted circle is the emerging ParC binding site and the amino acids involved in binding list in the below. (**C**) ParC docking on the new ParC binding sites of ParA dimer with mutation in C2.

**S7 Fig**. Homology model of ParA with ATP/ADP. Soj with ATP/ADP (6iub/6iud) [41] were used as templates. (**A**) Analysis of the interaction between ParA and ParC. (**B**) ParC assembled at C1 site of ParA-ATP. (**C**) ParC assembled at C2 site of ParA-ATP.

**S8 Fig**. (**A**) The *parC* gene and the corresponding ParC protein sequences, showing the sites for the in-frame deletion. The missing part was marked in red, the starting codon and amino acid were framed in green, the ending codon was framed in purple, and the ribosome binding sites was underlined in green. (**B**) Soluble expression analysis of ParC and ParA proteins in *E. coli* in different arrangements of *parC* and *parA*. The two genes were linked into the expression plasmid pET-15b according to the arrangement sketched in **Figure 1C**. (**C**) Protein aggregation analysis. ParA monomer was incubated at 32 °C and sampled at intervals (0, 5, 10 and 15 min), and the aggregation was mixed with 0.1% SDS and detected by native-page electrophoresis. Many tailed bands were observed in electrophoresis gel, probably result from different polymerization extents of ParA proteins.

**S9 Fig**. (**A**) A sketch for gene alternation of ParCA fusion protein (VII), and expression and purification of the ParCA fusion protein expressed in *E. coli* BL21(DE3) cells after IPTG induction. (**B**) Soluble expression of the ParC protein with the deletion of 14 C-terminal amino acids (plasmid XI). (**C**) Soluble expression of the ParCA fusion proteins with amino acid mutation at C2 site of the ParA domain (plasmid X). (**D**) Soluble expression of the ParCA fusion proteins with a 5×Ala linker (VIII) or an 8×Ala linker (IX) between the C and A domains. M, protein standard; S, supernatant; IB, inclusion body; FT, flowthrough; W, washing with 20 mM imidazole; E, elution.

## Supplementary tables

**S1 Table**. Function annotation of the pMF1 genes in GenBank and the homologous gene sources with the highest similarity.

**S2 Table**. Bacterial strains and plasmids used in this study.

**S3 Table**. Primers used in this study.

